# Extracting extended vocal units from two neighborhoods in the embedding plane

**DOI:** 10.1101/2022.09.26.509501

**Authors:** Corinna Lorenz, Xinyu Hao, Tomas Tomka, Linus Rüttimann, Richard H.R. Hahnloser

## Abstract

Annotating and proofreading data sets of complex natural behaviors are tedious tasks because instances of a given behavior need to be correctly segmented from background noise and must be classified with minimal false positive error rate. Low-dimensional embeddings have proven very useful for this task because they provide a visually appealing overview of a data set in which relevant clusters appear spontaneously. However, low-dimensional embeddings introduce errors because they fail to preserve high dimensional distances; and embeddings represent only objects of fixed dimensionality, which conflicts with natural objects such as vocalizations that have variable dimensions stemming from their variable durations. To mitigate these issues, we introduce a semi-supervised method for simultaneous segmentation and clustering of vocalizations. We define vocal units of a given type in terms of two density-based regions in low-dimensional embedding space, one associated with onsets and the other with offsets. We demonstrate our approach on the task of clustering adult zebra finch vocalizations embedded into the 2d plane with UMAP. We show that two-neighborhood (2N) extraction allows the identification of short and long vocal renditions from continuous data streams without initially committing to a particular segmentation of the data. Also, 2N vocal extraction achieves much lower false positive error rate than approaches based on a single defining region.

## 1 Introduction

One of the main problems in unsupervised data analysis is to rapidly obtain an overview of complex data without missing any possibly important details. For example, in the case of animal behavior, an investigator may want to rapidly inspect all the different types of vocalizations emitted by an animal. For this type of problem, dimensionality reduction techniques come in handy because they allow to display even high-dimensional data points such as complex vocal utterances on a two-dimensional computer screen (Sainburg et al. 2019; Sainburg et al. 2020; Kollmorgen et al. 2020). However, in distance-preserving embeddings such as TSNE (Maaten and Hinton 2008) or UMAP (McInnes et al. 2018), the distance between two points in the 2d-plane only approximates the true distance between the pair of vocalizations in the higher-dimensional space of the original data (Kollmorgen et al. 2020). In fact, embedding distances are not perfectly preserved because local neighborhoods in two dimensions are much smaller than the true neighborhoods in the high-dimensional space. How to efficiently deal with such embedding distortions remains a bottleneck in data browsing, proofreading, and annotation tasks (Chari et al. 2021).

Moreover, natural vocalizations tend to have variable durations, which clashes with the rigid dimensionality of embeddings. Although there are workaround techniques such as zero padding, these depend on segmenting the signal into foreground and background as a preprocessing step, which tends to introduce errors on its own and so may act against end-to-end extraction of vocalizations from raw data. All these caveats and challenges limit the widespread adoption of dimensionality reduction techniques for annotating and proofreading of vocalizations.

In general terms, the goal of our data annotation task is to extract flexibly defined and variably sized events from a continuous data stream. Our approach to vocal clustering is somewhat orthogonal to previous approaches where automated classifiers are optimized for the assignment of pre-computed segments to labels, either in a supervised (Tachibana et al. 2014; Nicholson 2016; Cohen et al. 2022) or unsupervised manner (Burkett et al. 2015; Morita et al. 2021). The goal of these approaches is to identify an efficient workflow that minimizes human involvement. In contrast, our aim is to place the human expert in in the center of the process, allowing him/her to explore and navigate, segment, and cluster recorded data in a fast and intuitive way using an interactive visualization and annotation tool.

We address the problem of clustering zebra finch vocalizations, which is to distinguish the diverse vocalizations produced by birds from all other sounds in the environment. In essence, the task is to correctly identify in midst of diverse noises all vocal units including song syllables and calls together with their onsets and offsets and their types. To robustly extract variable vocalizations from possibly distorted embeddings, we introduce for each vocalization type a pair of distinguishing feature sets, one anchored to the onset of the vocalization and the other to the offset. Because we extract vocalizations without segmentation as a pre-processing step, the specification of onset- and offset-anchored feature sets implicitly solves the segmentation problem.

## 2 Results

The first step of our vocal extraction method is to detect sound intervals (as opposed to intervals of silence) and to densely dissect the associated sound spectrograms into overlapping snippets, Figure 1. By considering each snippet as a potential vocalization onset or offset, the vocal segmentation problem remains unresolved at this step, it will be addressed at a later processing step.

**Figure 1:**
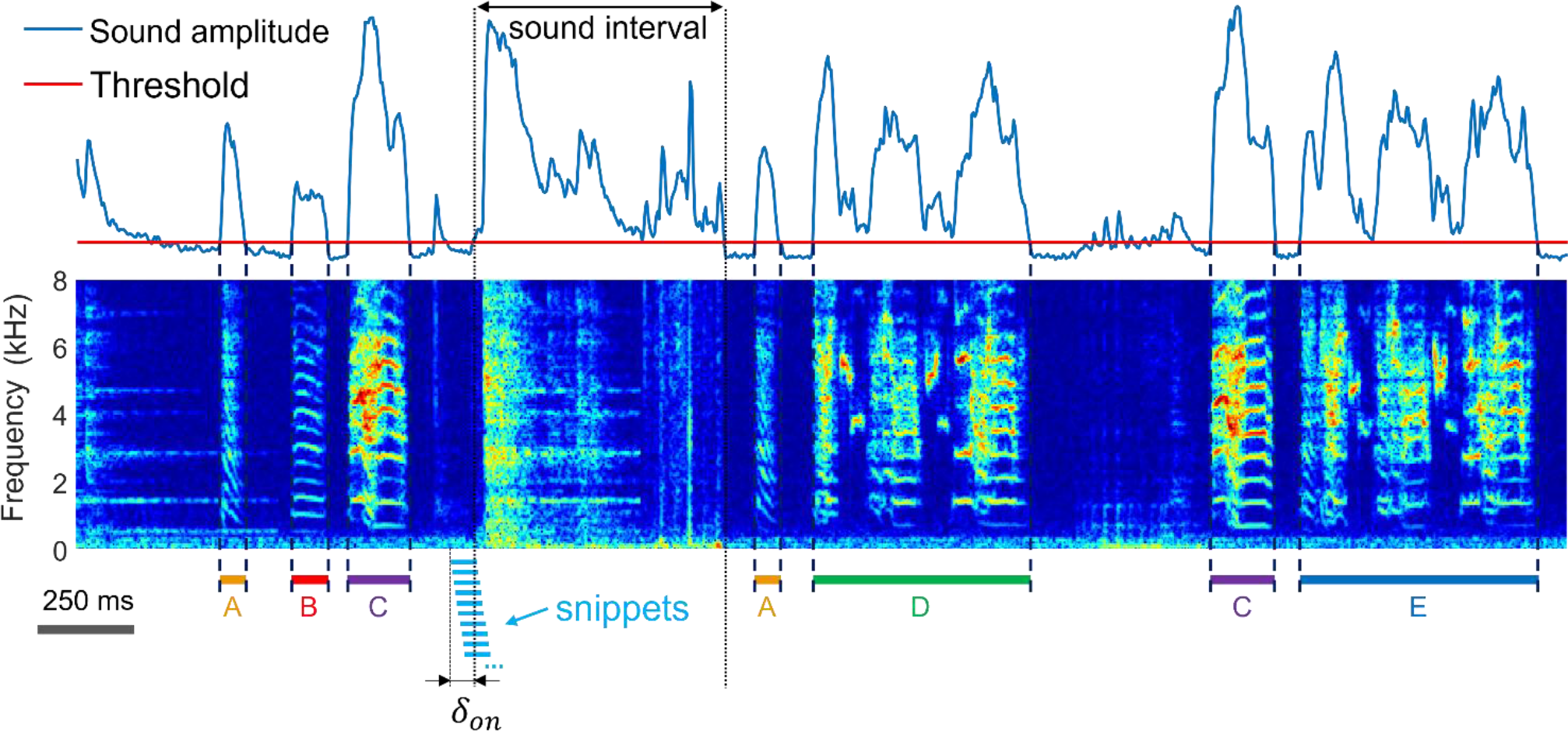
The vocal extraction problem is to correctly detect the onsets and offsets (dashed lines) of vocalizations and to determine their type (here A to D), amidst diverse non-vocal sounds (noise). Shown is a time-frequency log spectrogram of adult zebra finch song. Our approach to vocal extraction is to first extract sound intervals (based on threshold crossings of sound amplitude) and to dissect these into 64-ms long snippets (light blue bars) with 60 ms overlap among adjacent snippets. To achieve robustness to segmentation errors, we consider each snippet a potential onset or offset of a vocalization. The first snippet associated with an amplitude threshold crossing precedes the threshold crossing by a small margin δ_on_ (free parameter). The shown spectrogram was produced by concatenating diverse recording segments, chosen to illustrate the diversity of vocalizations and noises.

We then project all spectrogram snippets into the 2d plane, similar to the continuous UMAP embedding in Sainburg et al (2020). We assume that very often, a vocalization onset corresponds to a lower (low-to-high) threshold crossing of sound amplitude, and an offset corresponds to an upper (high-to-low) threshold crossing. Provided that for a given vocalization type, there are sufficiently many vocal renditions that are precisely segmented by sound amplitude (Fig. 1), the corresponding snippet embeddings will lie close to each other in the embedding plane and form a dense cloud of points. In general, we expect to find an extended cloud of 2d points that ranges from self-similar onset snippets on one end to self-similar offset snippets on the other.

When data is somewhat noisy, among the onset cloud of points, we will also find points that display a nonzero lag to the nearest threshold crossing of sound amplitude. In other words, in noisy data, we expect that not all vocalizations are cleanly segmented by sound amplitude but that instead some vocalizations are preceded or followed by a suprathreshold noise, which means that the time lag to an amplitude threshold crossing can be arbitrary large. Such noisy vocalizations can nevertheless be correctly extracted with our method because for extraction we rely not on prior segmentation but on similarity with cleanly segmented renditions.

We illustrate our extraction method on a one-day-long recording of an isolated male zebra finch. We sliced the recorded sounds into more than half a million snippets of 64 ms duration each that we embedded into the 2d place using UMAP, Figure 2.

**Figure 2:**
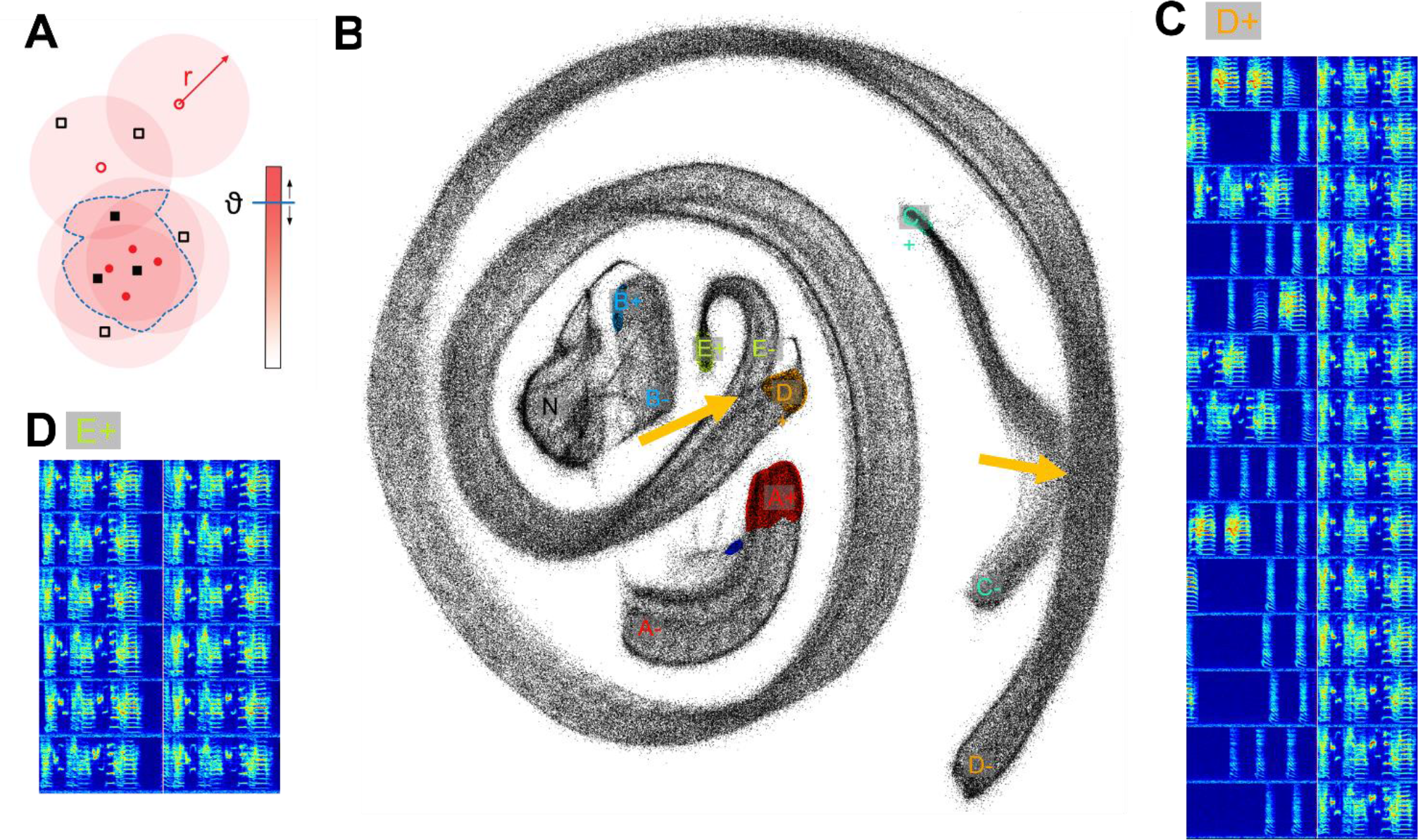
Two-neighborhood (2N) extraction of vocalizations from dense spectrogram embeddings. **A)** Schematic illustrating the definition of a blob. After projecting all data snippets into the 2D plane using UMAP, we replace the points corresponding to upward threshold crossings (d = 0, red circles, first light blue bar in Figure 1) by large, filled disks (red) of radius r that we sum up. All pixels at which the sum exceeds a given threshold ϑ (blue horizontal bar) are grouped into a blob (delimited by blue dashed line). All points that fall into this blob correspond to extracted vocalization onsets, including points but that were dissected at a lag different from d (black filled squares). **B)** Projected data snippets (black dots) from a one-day long recording. The onset blobs corresponding to the time slice d = 0 are shown in color (different colors for different vocalization types). The yellow arrows point to indistinguishable snippet embeddings stemming from different vocalization types. The letters A-E indicate manually chosen onset slices ‘+’ and offset slices ‘-’ for each vocalization type (same lettering as in Figure 1). The cluster labelled ‘N’ (gray) is a noise cluster without distinct onset an offset behavior, this cluster was ignored. The small blue blob next to the A+ blob is an onset variant of the introductory note, which can either be included in the definition of A+ (introductory note) or excluded. **C), D)** Spectrograms of example syllables taken from blob D+ (C) and blob E+ (D). Same bird as in Figure 1.

In practice, for each vocalization type, we define at least one dense region of points (a blob, see Figure 2) in the 2d plane associated with the onsets of this vocalization type, and at least one blob with the offsets. The extracted individual vocalizations are then defined as the spectrogram chunks delimited by the timestamp of a point in an onset blob and the timestamp of the first subsequent point within an offset blob, i.e., we extract vocalizations simply within the shortest time intervals between pairs of points in onset- and offset blobs. This method is oblivious of the time lags to amplitude threshold crossings of the underlying points, and so our method can correctly extract vocalizations that are not cleanly segmented by sound amplitude.

In more detail, our vocalization extraction method consists of choosing onset/offset blobs defined via a manually chosen time lag *d* relative to the adjacent amplitude threshold crossing and to harvest all points within the blob, including points that were sliced at different time lags (see Methods). Typically, *d* = 0, but the lag variable *d* provides a degree of freedom that we use to minimize effects of embedding distortions and to disambiguate confounding vocalization types, as detailed in the following.

Note that snippets from different syllables can appear indistinguishable on the 2D plane, Figure 2, either because a bird repeats a sub-syllable or note in a different context (the two are truly indistinguishable) or because of UMAP projection errors (the latter we visually found to be quite common). When such non-discriminability occurs at either an onset or an offset, the respective snippet then loses its distinguishing characteristic for that syllable, Figure 2C. For example, zebra finch song has the characteristic that birds tend to sing different syllable types with indistinguishable endings. This ambiguity implies that the spectrogram snippets near an upper threshold crossing do not uniquely define the ending of that syllable type (presumably the same is also true for some syllable onsets). As a workaround to such repetitive structure inherent in birdsong (and language for that sake), our method provides the freedom to define syllable endings and beginnings at fixed time lags away from threshold crossings, at places within a syllable where the defining snippet becomes unique for that syllable. Using our GUI (Supplementary Material), users can increase and decrease the lag variable *d* to observe the blobs move around in the embedding plane until they reach a region in the 2d plane where there are no confounding points from other vocalization types. Such confounding points can be recognized thanks to the elongated 1d-structure that densely dissected vocalizations obtain in the 2d plane (Sainburg et al. 2020): the confounding points are the ones where two different 1d-structures come too close to each other (see Fig. 2B, yellow arrows). As can be seen in Figure 2B, in the chosen bird, the endings of syllables D and E coincided with each other in the embedding plane, which is why we had to define the offset-anchored blob D-not far from the onset-anchored blob D+ in a region where it was distinct from any neighborhood of E-, to make sure the endings of syllable D are not confounded with parts of syllable E. Namely, this bird produced two very long and complex song syllable types in rapid succession, whereby the second type (E) displayed an additional small down sweep at the syllable beginning, making it clearly distinct from Syllable D only by virtue of this down sweep, Figure 2C, D. In addition to the time lag *d*, the definition of a blob also depends on two parameters: a radius *r* and a threshold *ϑ* that set the size of a blob as a function of the local density of points. In essence, the radius *r* sets the sizes of the disks that are placed at the locations of the embedding points and the threshold *ϑ* is the height that the summed disks must exceed for a pixel to be included in a blob, Figure 2A.

When extracting vocalizations, we perform the following steps:

1. Set a radius *r* and threshold *ϑ* to result in a visually appealing blob that is well matched to the density and scattering of the snippet embeddings for the vocalization type of interest.
2. Change the time lag variable *d* to a certain positive (negative) value relative to the lower (upper) threshold crossing and observe the blob moving along the data.
3. Place the blob in a place far away from all other vocalizations to define a unique neighborhood anchored to the vocalization onset (offset). The onsets (offsets) of renditions of this vocalization type are given by the timestamp of all snippets falling within that blob minus the selected time lag *d*.
4. Possibly add more blobs to the definition of the onset (offset) of the vocalization type of interest. Namely, defining regions of an onset (offset) of a given vocalization type can be formed by a non-connected set of blobs, each at a distinct time lag.

Once both onsets and offsets are defined, we extract as renditions of this vocalization type the spectrogram regions delimited by adjacent onsets and offsets. For the bird shown in Figure 1 and 2, about 96.6% of thus-extracted vocalizations had cleanly segmented onsets and about 98.7% had cleanly segmented offsets (i.e. onsets and offsets coincided with sound amplitude threshold crossings). In another bird, also recorded with a microphone and kept alone in a soundproof box, clean segmentation was even more frequent (99.9% for onsets and 99.0% for offsets, respectively). However, in birds housed in pairs and recorded with a wireless accelerometer mounted to their back as in Anisimov et al (2014), the fraction of cleanly segmented vocalizations was much lower (down to 72% for onsets and 89% for offsets, respectively). Thus, the level of noise depends strongly on the recording method, but our method allows harvesting vocalizations even in noisy situations.

How good are the extracted vocalizations? We computed two performance measures associated with the extraction procedure: 1) The *quality of the segmentation* in terms of the time differences of extracted onsets and offsets relative to gold standard human annotations; and 2) the *clustering performance* in terms of the false-positive error rate of misclassified time bins, again assessed by human experts.

With regards to the quality of the segmentation, for the bird shown in Figures 1 and 2 and for both onsets and offsets, the time lags to threshold crossings were similarly distributed for 2N- and for human-extracted vocalizations (Fig. 3A, B), suggesting that 2N extraction is able to extract vocalizations from background noise in similar manners as humans do. For both onsets and offsets in this bird, the largest time lags to amplitude-threshold crossings were up to 1 s long, revealing a payoff for getting rid of segmentation as a pre-processing step. In all birds tested (n=2 mic and n=2 accelerometer birds) we found similarity between 2N-extracted vocal segments and the human gold-standard counterparts, constituting a big improvement over simple sound amplitude thresholding (Fig. 3C, D).

**Figure 3:**
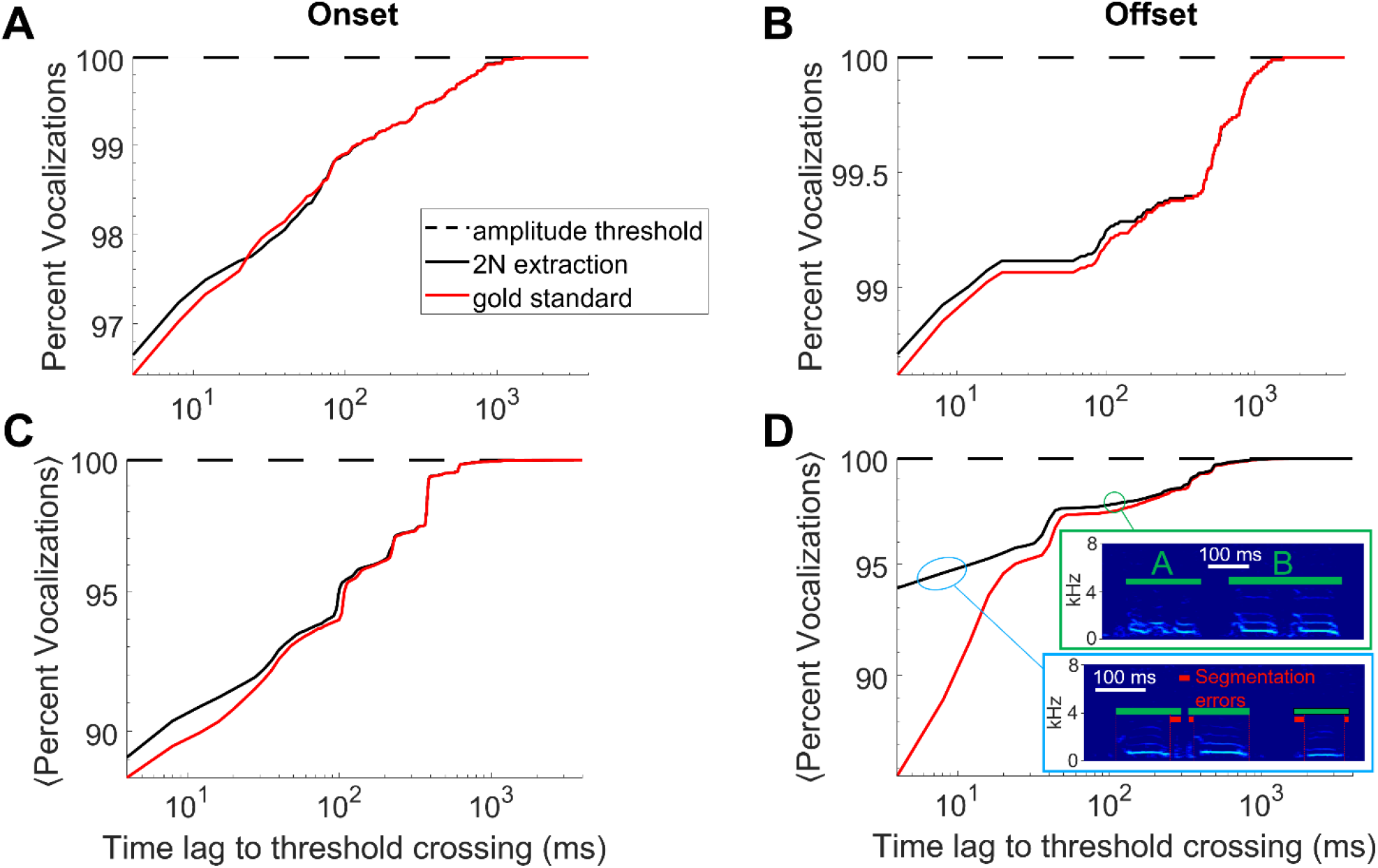
2N-extracted vocalizations (black curve) are similarly segmented as human-extracted vocalizations (red). **A)** Shown is the cumulative percentage of vocalizations with an onset that falls within a given time lag (x-axis) following a sound amplitude threshold crossing. **B)** Same for offsets that are within a given time lag preceding a sound-amplitude threshold crossing. Same bird as in Figure 1. **C,D)** Cumulative percentage averaged across n=4 birds. The insets in D show typical errors made by 2N extraction (green bars), which is to interpret a string of two calls as a single call (here labelled ‘B’, green bounding box) or to introduce segmentation errors (red bars) from too generous inclusion of surrounding noises (blue bounding box), data taken from an accelerometer-recorded bird. (A-D) Segments extracted by amplitude threshold crossings (dashed) trivially display no time lag whatsoever.

Some disagreements were seen when data was noisy; namely, we observed a tendency in human evaluators to segment the offsets earlier, which we found was often due to double misses that occurred in strings of calls where both an offset and the following onset were missed, leading to the hallucination of a much longer call than there actually was, Figure 3D. We do not propose a workaround for such problems here, but these problems are fixable in practice by detecting for each vocalization type the outlier renditions of excessively long duration and by ignoring these. The shorter segmentation errors within 10-20 ms of true syllable offsets were almost always due to inclusion of respiratory or movement artifacts near syllable boundaries, Figure 3D, and very rarely were they due to truncations of parts of syllables, which would be more detrimental for subsequent feature-based syllable analysis. In summary, our method provides an improved vocal segmentation compared to simple sound amplitude thresholding, in particular when recordings are noisy such as when birds are pair-housed and recorded with animal-borne sensors.

We assessed vocal clustering performance of 2N extraction by asking an expert to fix all extraction errors, which in addition to the segmentation errors were the classification errors of either assigning a noise to a vocalization or a vocalization to the wrong type. More specifically, we evaluated the precision of 2N extraction in terms of the fraction of 4-ms time bins that were assigned to the correct vocalization type as determined by a human expert. We were particularly interested in comparing our findings to a baseline of extracting vocalizations not from two defining (sets of) regions in the embedding plane, but only from a single region, either anchored to the onset or the offset, but not both. In these 1N extraction baselines, we extracted vocalizations from a point in a blob until either the next point in the blob or until the end of the sound interval, whichever came first (see Methods).

We found that 2N extraction outperformed 1N extractions by a large margin, achieving 3-6 times lower extraction error, Figure 4. The superior precision of 2N extraction came only at a minimal cost of lower recall. Namely, for the bird shown in Figure 1, 2N extraction retrieved almost as many time bins as did 1N extraction, namely 99.6%. On average (n=4 birds), the number of time bins retrieved with 2N extraction was 96.9% relative to the mean number of bins retrieved with 1N extractions (averages across onset-and offset-based 1N extraction methods). Thus, the added benefit of much lower extraction error came only at a minimal cost of potentially retrieving fewer vocalizations. Thus, in terms of extraction performance, it pays off to extract vocal units in terms of two sets of defining characteristics, one anchored to the onset and the other to the offset. In terms of manual processing time, 2N extraction comes at the obvious cost of twice the workload compared to more traditional 1N extraction. However, given that 2N extraction can be routinely done within less than five minutes for an experienced user, irrespective of the amount of data that needs to be processed, this overhead seems negligible in practice.

**Figure 4:**
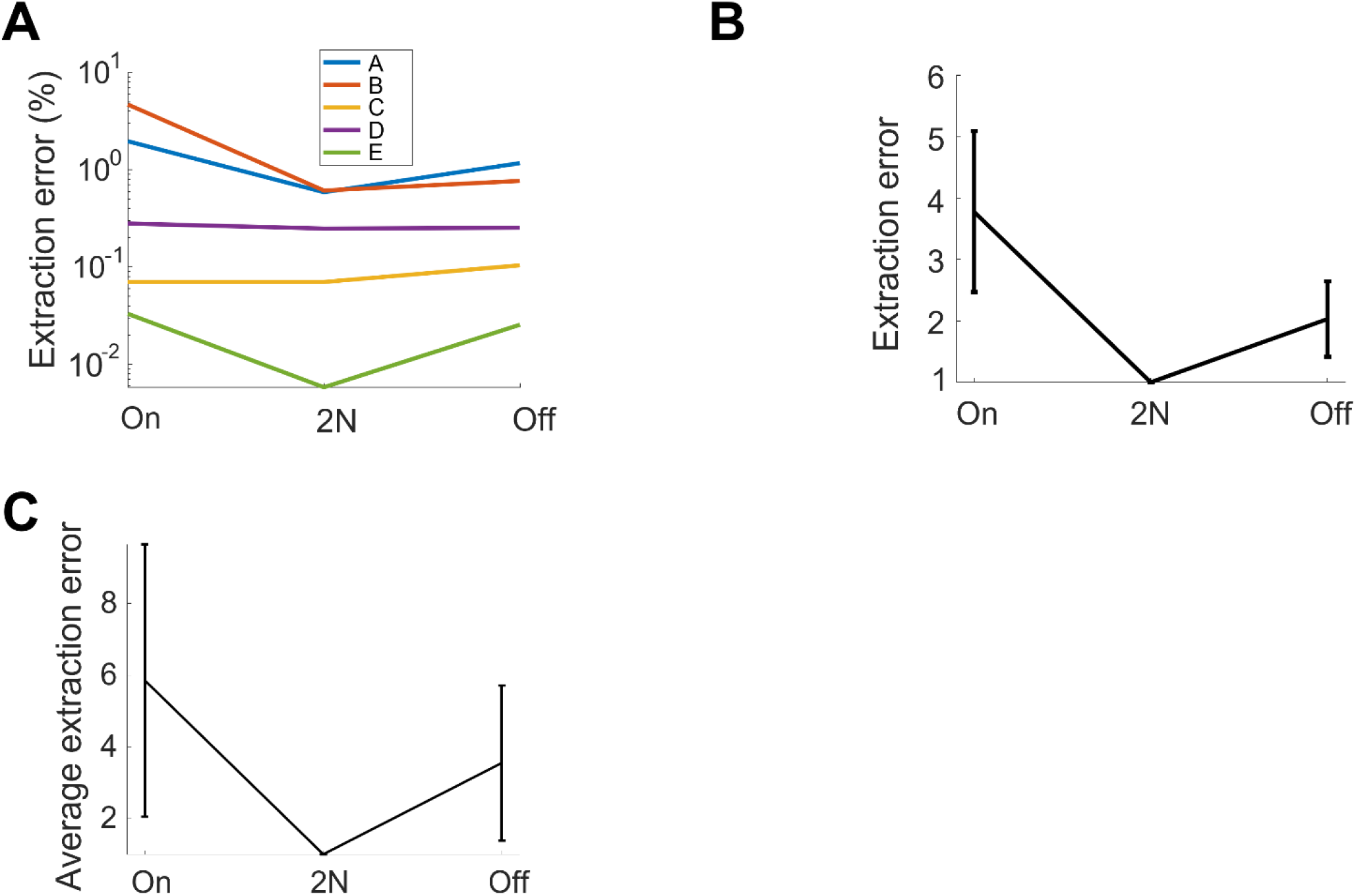
2N-extracted vocalizations achieve higher precision than their 1N-extracted counterparts. The fraction of extracted (4-ms) time bins that are mis-classified (extraction errors) is shown for diverse methods: 2N extraction and 1N extraction, either onset-anchored (on) or offset-anchored (off). **A)** For each vocalization type, the error is lower when vocalizations are extracted from two neighborhoods than when extracted from one neighborhood. Same bird as in Figure 1 and 2. **B)** Same data, averaged over all syllable types and after normalizing by the error rate of 2N extraction (shown is average ± std across vocalization types). The retrieval error of 1N extraction is 2-4 times higher than that of 2N extraction. **C)** Normalized relative extraction error (average ± std across 4 birds). 2N extraction achieves 3-6 times fewer errors.

## 3 Discussion

We presented a simple and intuitive method for extracting arbitrary vocal units from a continuous stream of data. Our key contribution is to identify a particular vocalization not through a single set of characteristics tied to a single component such as the onset, the offset, or even an entire vocalization, but through two sets of characteristics, tied to two components near the onset and offset. The benefits of this dual recognition are better segmentation and higher clustering performance, because errors resulting from vocal ambiguities and embedding distortions will less likely happen near both the onset and offset of a given vocal instance rather than just either one of them. Accordingly, our method works best when onset and offset defining blobs are sufficiently far apart from each other such that there is no overlap between them.

2N extraction makes most sense on large data sets because there is only a time penalty for computing the embedding but virtually no overhead for defining the onset and offset blobs. We routinely calculated UMAP embeddings of up to 1 million sound snippets using standard desktop PCs with 32 GB of RAM. Embedding methods such as UMAP and t-SNE have been criticized for the distortions they can create, especially in genomic data (Chari et al. 2021). While we find similar distortions in vocal data, our workaround is to flexibly define vocal units based on regions in the embedding plane that are far from ambiguities and presumably also from distortions. But our method is not tied to a particular embedding method and we expect it to work also on other 2d embedding types.

2N extraction is very flexible. As noted, the defining regions of a vocalization type need not constitute a connected set. For example, we could have chosen to combine syllables D and E in Figure 1 into a single syllable type by defining its onset characteristic in terms of the two blobs labeled D+ and E+ in Figure 2. Thus, our GUI provides the user with high flexibility of defining vocal units, which minimizes the need for postprocessing including the correction of segmentation errors.

According to our parameter choice, the first and last snippets protrude by *δ*_*on*_ and *δ*_*off*_ beyond the underlying sound intervals, which provides the main advantages that fewer data points need to be embedded (providing a speed-up) and that vocalizations tend to appear as distinct 1D structures with clear boundaries rather than as a single excessively long 1d structure. Currently, our method requires a pre-segmentation into sound intervals, as otherwise we do not obtain a time lag *d* that we need to define the blobs and to extract vocal units. In data that is so noisy that there are barely any vocalizations that are cleanly segmented by sound amplitude, our method is not trivially applicable. However, in this case, it may be possible to use another sound feature than sound amplitude, as long as the feature is able to identically dissect a significant number of vocalizations and provided these appear as a compact cluster in the embedding plane such that a blob will emerge using our procedure in Figure 2A. We imagine that our approach to signal extraction can generalize to biological and physical processes other than vocalizations. Namely, we believe that our approach will work well for the extraction of units in natural processes that contain rigid elements that sequentially unfold in variable sequences and at variable speeds. The duration of objects of interest should be typically longer than the snippet size. When overlapping data snippets from such processes are projected onto the 2D plane, elongated structures will result, ideally displaying uniquely defining beginnings and endings as in Figure 2B. The method will likely also work for spatial rather than temporal data, provided that the same requirements of repetitive sequential structure applies. We hope that our GUI can be of use to researchers wanting to adopt our methods for their work and as a basis for further developments.

## 4 Methods

Our data stem from single-housed birds recorded with wall-mounted microphones or from pairs of birds which wear harnesses carrying accelerometers that signal body vibrations stemming from self-produced vocalizations. Although all birds have been recorded in acoustically isolated environments, the vocal extraction problem is challenging in species such as the zebra finch that produce not just harmonic sounds but that also emit broadband vocalizations which can resemble non-vocal sounds in their cage.

We sampled signals at 32 kHz and computed log-power sound spectrograms using the short-time Fourier transform in 512-sample windows and hop size among adjacent windows of 128 samples (i.e., 4 ms). We pre-segmented the data into sound intervals (assuming that without sound there is no vocalization) by thresholding the spectral power of microphone signals in the range 312 Hz to 8 kHz and of accelerometer signals (illustrated in Figure 3D) in the range 312 Hz to roughly 4 kHz. The threshold for sound interval extraction (Figure 1) was set to 5 standard deviations above the average spectral power calculated during periods of silence.

2N extraction was performed by densely dividing the spectrograms (within sound intervals) into spectrogram snippets of fixed width in the range 12-16 columns (corresponding to a duration of 48-64 ms). The hop size between adjacent segments was given by one column (i.e, 4 ms), Fig. 1. To both the first and last data snippets associated with a sound interval, we ascribed an integer time lag *d* to sound onset/offset of *d* = 0 (Fig. 2). The onset time *δ*_*on*_ of the first snippet preceded the sound interval onset by *δ*_*on*_ = 4 spectrogram columns (i.e. 16 ms, Fig. 1). Similarly, the offset time of the last snippet exceeded the sound interval offset by *δ*_*off*_ = 6 (i.e. 24 ms). These choices ensured that brief silent gaps before and after syllables were included in the defining characteristics of a vocalization. By definition, the second snippet of sound interval had a time lag of *d* = 1 (i.e. 4 ms) and the second-last snippet had a time lag of *d* = −1 (i.e. -4 ms), etc. This dissection of the data into snippets resulted in a total of 581k snippets for the day-long recording of the bird shown in Figure 2 (given the chosen snippet duration of 64 ms).

We then embedded the snippets into the 2d plane using UMAP (McInnes et al. 2018) and created an image by replacing all points in the embedding plane associated with a given positive integer lag *d* to sound onset (or negative integer lag to sound offset) by a small disk of radius *r*. In fact, to obtain larger blobs, we also replaced points in adjacent snippets (at lag *d* + 1 for onsets and lag *d* − 1 for offsets) by such disks. Summing these disks and thresholding the resulting sum with a threshold *ϑ* resulted in the blobs shown in Figure 2.

By changing the lag value *d*, blobs were interactively moved to uniquely defining regions along the 1d manifold of a vocalization type where no points from other vocalizations could be found. All 2d points falling into a thus-placed onset blob were then associated with that vocalization type, with vocalization onset times given by the chosen snippet timestamp (i.e. the earliest snippet in case several adjacent points were found) minus the chosen time lag *d* (the smaller lag in case blobs were defined by two time slices). The time differences between the extracted vocalization onsets and the start times of the underlying sound intervals are shown as a cumulative density in Figure 3A, the corresponding density for vocalization offsets relative to sound-interval endings is shown in Figure 3B.

To estimate the precision of 2N extraction, we determined for all 2N-extracted vocalizations of a given type, the fraction *f* of correctly classified 4-ms time bins, as assessed by a human expert. We then repeated the same estimates for 1N-extracted vocalizations that were defined by considering only the onsets blobs or only the offset blobs rather than both. That is, in the onset-anchored 1N baseline, we extracted a vocalization from the time difference Δ*t* = *t*_*a*_ − *t*_*b*_, where *t*_*b*_ is the timestamp of a point in the onset blob minus the selected time lag *d* and *t*_*a*_ is the end of the underlying sound interval or the timestamp of the next point in the onset blob, whichever comes first. Analogously, in the 1N offset baseline, we extracted a vocalization from the timestamp of a given point in the offset blob minus the selected (negative) time lag *d*, backwards, until the previous point in the offset blob or the beginning of the underlying sound interval, whichever comes first (going backwards in time). The corresponding extraction errors *ϵ* = 1 − *f* are shown in Figure 4 for diverse vocalization types and birds.

### 4.1 2N extraction algorithm

In the following, we describe the detailed extraction procedure of a vocalization, starting with the onset. Extraction is parameterized by three variables: a time lag *d*, a radius *r*, and a density threshold *ϑ*. We perform the following steps:

1. Define an integer time lag *d* ≥ 0 as small as possible (start with *d* = 0).
2. Identify all 2d points associated with this lag *d*.
3. Replace each identified point with a disc of radius *r* and sum up these discs, yielding a 2d density.
4. Identify the regions where the 2d-density exceeds a threshold *ϑ*. These regions we refer to as blobs, they are the defining characteristics of vocalization onsets.
5. Change *d, r*, and *θ* to place and shape the blob such that it defines a uniquely characteristic region of the vocalization of interest. The time lag *d* is the length of the anchor associated with that blob.
6. Repeat steps 1-5 up to *K* times to define diverse onset blobs for a given vocalization type (typically, *K* = 1 because there is a unique blob for each vocalization type).
7. Identify all points inside the *K* blobs. These points uniquely define the onsets of the extracted vocalizations given by their timestamps minus the anchoring time lag *d*_*i*_ of the underlying blob (*i* = 1, …, *K*).

In this procedure, the optimal choices of radius *r* and threshold *ϑ* depend on the local density of points and should be individually chosen for each vocalization type and each blob. Essentially, the radius should be chosen as large as possible to not miss any onsets and likewise the threshold should be as low as possible to make the blob as large as possible to maximize the point harvest. However, the blob should not extend into points associated with confounding vocalizations, so a bit of manual fine-tuning is required for each vocalization type.

When all onsets of a vocalization type are extracted, we perform the analogous extraction of offsets; the lags of offset-defining blobs satisfy *d* ≤ 0; the parameters *d, r, ϑ*, and *K* of offset blobs need similar fine-tuning. We first defined the onset and offset blobs of all vocalizations and only then we extracted the vocalizations.

Based on all defined blobs, we then extracted the vocalizations themselves as the spectrogram regions delimited by an onset and a subsequent offset. Cases in which an onset was followed by an offset of another type were discarded. Also discarded were onsets following an onset with missing intervening offset (and vice versa offsets following an offset with missing intervening onset). The resulting selections were visually inspected and manually corrected in terms of segmentation and clustering errors. That is, onset and offset times were manually adjusted by an expert in 4 ms steps to the nearest true onset or offset; and, misclassified vocalizations were assigned to the correct vocalization type.

### 4.2 Practical notes

In all adult zebra finches examined, we managed for each vocalization type to find distinct onset and offset blobs. We carefully selected each blob as close as possible to its extreme position, i.e., as close as possible to the onset resp. offset of the associated sound interval. In nearly all birds, we found at least one cloud of 2d points corresponding to non-vocal noise; in this cloud it was impossible to select both onset and offset blobs, presumably because noise tends to be unstructured, i.e., for noise there were no distinct time lags to amplitude threshold crossings at which noise snippets appeared more similar with each other than with snippets at other lags. We therefore extracted noise segments as the time intervals from the extracted onset until the ending of the underlying sound interval or the next vocalization onset, whichever came first.

## 6 Author Contribution

RHRH, XH, TT, LR and CL contributed to the concept and design of the study. LR recorded data from backpack experiments. RHRH implemented the algorithm and initial GUI and performed the analysis. RHRH and CL created visualizations and wrote the manuscript. CL refined the GUI. XH, TT, and LR edited and provided feedback on early versions of the manuscript and the GUI.

## 7 Acknowledgement

We thank Anja Zai for performing additional song recording experiments and Heiko Hörster for his help thereof. All experimental procedures were approved by the Veterinary Office of the Canton of Zurich.

